# Establishing combination PAC-1 and TRAIL regimens for treating ovarian cancer based on patient-specific pharmacokinetic profiles using *in silico* clinical trials

**DOI:** 10.1101/2022.03.29.486309

**Authors:** Olivia Cardinal, Chloé Burlot, Yangxin Fu, Powel Crosley, Mary Hitt, Morgan Craig, Adrianne L. Jenner

## Abstract

Ovarian cancer is commonly diagnosed in its late stages, and new treatment modalities are needed to improve patient outcomes and survival. We have recently established the synergistic effects of combination tumour necrosis factor-related apoptosis-inducing ligand (TRAIL) and procaspase activating compound (PAC-1) therapies in granulosa cell tumours (GCT) of the ovary, a rare form of ovarian cancer, using a mathematical model of the effects of both drugs in a GCT cell line. Here, to understand the mechanisms of combined TRAIL and PAC-1 therapy, study the viability of this treatment strategy, and accelerate preclinical translation, we leveraged our mathematical model in combination with population pharmacokinetics (PopPK) models of both TRAIL and PAC-1 to expand a realistic heterogeneous cohort of virtual patients and optimize treatment schedules. Using this approach, we investigated treatment responses in this virtual cohort and determined optimal therapeutic schedules based on patient-specific pharmacokinetic characteristics. Our results showed that schedules with high initial doses of PAC-1 were required for therapeutic efficacy. Further analysis of individualized regimens revealed two distinct groups of virtual patients within our cohort: one with high PAC-1 elimination, and one with normal PAC-1 elimination. In the high elimination group, high weekly doses of both PAC-1 and TRAIL were necessary for therapeutic efficacy, however virtual patients in this group were predicted to have a worse prognosis when compared to those in the normal elimination group. Thus, PAC-1 pharmacokinetic characteristics, particularly clearance, can be used to identify patients most likely to respond to combined PAC-1 and TRAIL therapy. This work underlines the importance of quantitative approaches in preclinical oncology.

## Introduction

Ovarian cancer is one of the main causes of female cancer death^1^. Due to a lack of early diagnostics and the absence of early warning symptoms, patients with ovarian cancer are usually diagnosed at an advanced stage^1^. This late presentation is one of the main reasons for the high death rate of ovarian cancer. While curative treatment exists for patients with early stage disease, most of those who present with advanced disease will experience recurrences with progressively shorter disease-free intervals^2^. These episodes culminate in chemoresistance and ultimately bowel obstruction, the most frequent cause of death^2^. The current standardized treatment for ovarian cancer is optimal cytoreductive surgery plus platinum-based chemotherapy with carboplatin-paclitaxel^1^. However, the development of chemotherapy-resistant and refractory disease results in an overall reduction in sensitivity to chemotherapy and decreases to the long-term survival for ovarian cancer patients. As such, it is paramount that new targeted therapies for ovarian cancer are developed and investigated for their potential use in patients.

One of the main mechanisms of tumour cell killing by anti-cancer drugs is to induce programmed cell death, or apoptosis^3^. Procaspase activating compound-1 (PAC-1) is a small apoptosis activating molecule identified in a high-throughput screen of ~20,500 small-molecule compounds^4^. PAC-1 and its derivatives have since gone on to show efficacy as anti-cancer agents *in vitro*^3,5^ and *in vivo*^6,7^, and have demonstrated safety in animal models^5–7^. PAC-1 is also under investigation in phase I trials against advanced malignancies (NCT02355535, NCT03332355), and has the potential to become a novel and effective anticancer agent^8^. However, the ideal dose of PAC-1 in humans remains to be determined. Further, given heterogeneity in PAC-1 pharmacokinetics, the ability to attain effective plasma concentrations in patients is a challenge for clinical translation.

Tumour-necrosis factor (TNF)-related apoptosis-inducing ligand (TRAIL) is a transmembrane protein that triggers cellular apoptotic pathways. In cancer cells, TRAIL has been shown to induce apoptosis without toxicity to normal cells and is well-tolerated by patients^9,10^. Unfortunately, clinical trials with soluble TRAIL have failed to display efficacy^11^, which could be related to its short half-life *in vivo* and an inability to reach a therapeutic concentration at the tumour site^12^. As such, like PAC-1, to facilitate preclinical-to-clinical translation of TRAIL, it is imperative to gain an improved understanding of its pharmacokinetics.

We recently demonstrated the therapeutic efficacy *in vitro* of PAC-1 and TRAIL alone and in combination in granulosa cell tumours (GCT) *in vitro* models^3^. GCT constitute only ~5% of ovarian neoplasms, with up to 80% recurrent GCT resulting in fatality, motivating the search for more effective treatments against these tumours. Our results showed synergy between TRAIL and PAC-1, with this combination resulting in rapid kinetics of apoptosis induction and killing of GCT cell lines as well as patient samples of primary and recurrent GCT.

To facilitate clinical translation of combined PAC-1 and TRAIL in GCT, in this paper, we developed a mathematical model integrated with pharmacokinetics/pharmacodynamics (PK/PD) of both drugs. Taking the PK/PD models to be representative of an average patient, we sought to understand combination PAC-1/TRAIL therapy in a cohort of heterogeneous virtual patients using an *in silico* (virtual) clinical trial^13^. A virtual clinical trial^14^ is a complementary approach to clinical trials. In a virtual clinical trial, a heterogeneous population of *in silico* individuals is generated and an exploratory drug trial is conducted within that population to determine characteristics of responders/non-responders, determine mechanisms of drug action, and to optimize/personalize therapy^15^. Crucially, virtual clinical trials can also be used to suggest guidelines for inclusion in real clinical trials, helping to reduce attrition along the drug development pipeline. Virtual clinical trials have become increasingly popular in cancer drug development, as they enable optimal and robust therapeutic treatments to identified prior to human trials and at a lower cost to drug developers and patients^16,17^. Additionally, *in silico* clinical trials have proven to be useful in other areas of medicine where population heterogeneity is crucial to disease prognosis^18–20^.

To examine the efficacy of novel PAC-1 and TRAIL combination therapy while considering the impact of PK/PD heterogeneity, we extended our previous work^3^ to run a virtual clinical trial of this treatment. Using our previous model describing the cell viability under treatment with PAC-1 and TRAIL, we considered the impact of heterogeneity in PK parameters on optimal combined therapeutic regimens. In a generated population of 400 virtual patients, we deployed an *in silico* clinical trial approach similar to ones we have developed for combination oncolytic virotherapy^16,21^. We then individualized therapeutic regimens for each of the 400 virtual patients and determined characteristics of responders. Our results suggest that the PKs of PAC-1 determine both treatment regimens and efficacy, and may be used to identify patients who will most benefit from combination PAC-1 and TRAIL therapy. Together, this work highlights the use of quantitative modelling in preclinical cancer therapy development, and the potential of combined PAC-1 and TRAIL to treat GCT.

## Methods

### GCT cell count measurements

The human adult GCT cell line KGN (Riken BioResource Research Center, Ibaraki, Japan) was cultured in Dulbecco’s modified Eagle’s medium/nutrient mixture F12 (DMEM/F12, Sigma-Aldrich, St. Louis, MO, USA) with 5% fetal bovine serum (FBS) (Gibco, Waltham, MA, USA). The proliferation rate of KGN cells was assessed by first seeding into 24-well plates at 10,000 cells per well to ensure that cells remained subconfluent for the duration of the assay. Two technical replicates were counted daily for 7 days using a hemocytometer following trypsinization and trypan blue staining of the monolayers. The assay was repeated 3 times.

### Mathematical model of the combined effects of PAC-1 and TRAIL on ovarian tumour growth

To model the impact of dual drug combination therapy with PAC-1 and TRAIL, we previously developed a simple system of ordinary differential equations describing the temporal evolution of ovarian cancer cells (*G*(*t*)) and the effects of PAC-1 (*C*_*PAC1*_) and TRAIL (*C*_*TRAIL*_) concentrations^3^. We modelled tumour cell growth to follow a Gompertzian model, which is commonly used in cancer modelling to reflect saturating growth dynamics^22–25^. We considered simple exponential decay for the pharmacokinetics (PKs) of both PAC-1 and TRAIL, modelling each with linear elimination at rates *k*_*ePAC1*_ and *k*_*eTRAIL*_ respectively. The model is summarized in **Figure 1**A, and described by

**Figure 1.**
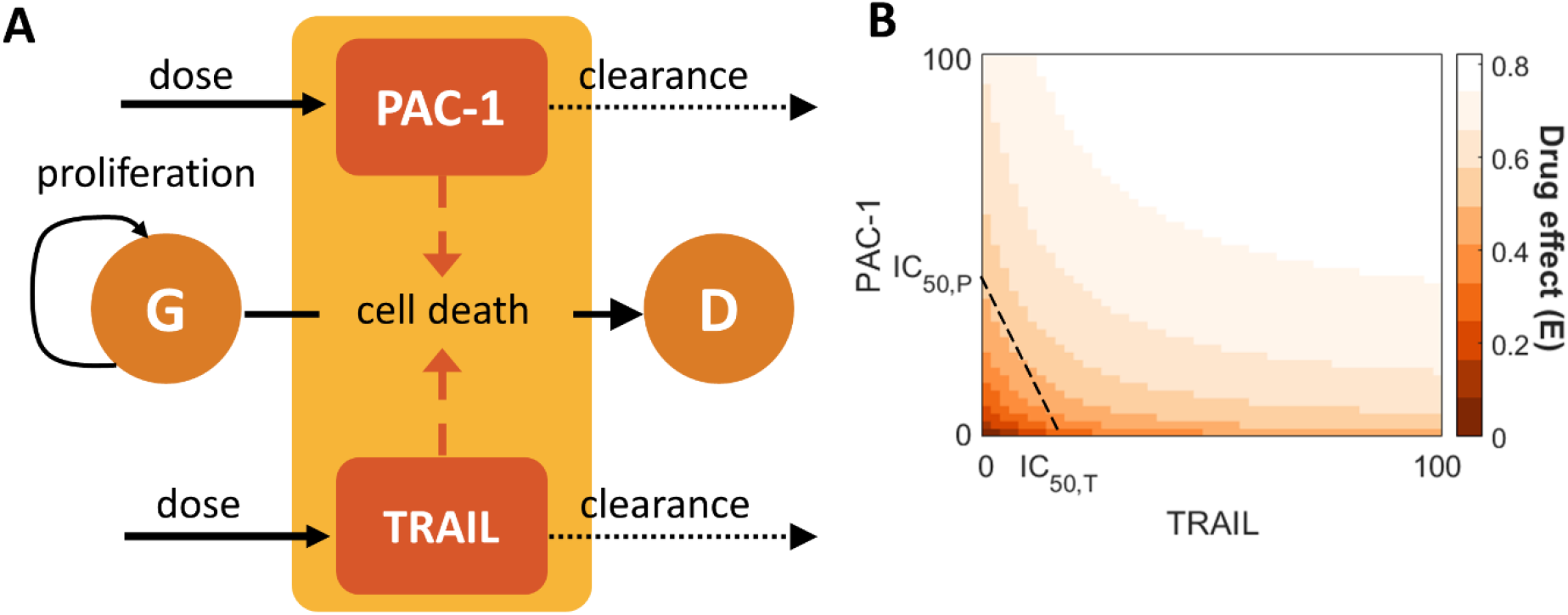
PK/PD model for synergistic effects of PAC-1 and TRAIL on GCT. (A) After administration, both drugs enter the blood stream and are subsequently linearly cleared. Tumour cells, *G*(*t*), proliferate according to Gompertzian growth. PAC-1, *C*_*PAC1*_(*t*), and TRAIL, *C*_*TRAIL*_(*t*), cause tumour cells death according to the combination effect function *E*(*C*_*PAC1*_(*t*), *C*_*TRAIL*_(*t*)) in **Eq. 4**. (B) The drug interaction profile for PAC-1 and TRAIL (**Eq. 4**) is plotted with their half-effect concentrations *IC*_50, *P*_ and *IC*_50, *T*_ noted. For given concentrations of PAC-1 and TRAIL denoted along the vertical and horizontal axes, respectively, the corresponding drug effect *E*(*C*_*PAC1*_(*t*), *C*_*TRAIL*_(*t*)) is shown (shaded orange regions). The dashed black line connecting the concentrations of both drugs achieving the same inhibitory effect is plotted to demonstrate the synergistic effect of both drugs, highlighted by comparing the concavity of the isoboles with the equidosage line joining the half-effect of either drugs^28^.

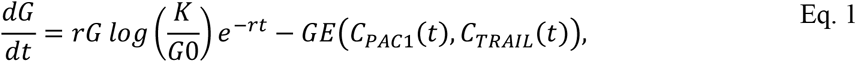

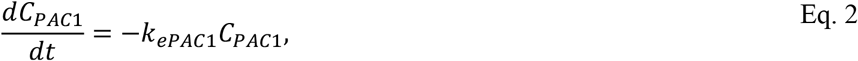

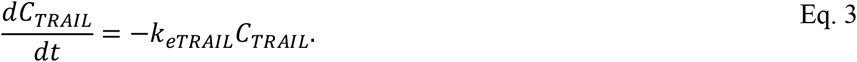

As in our previous work^3^, the combined (synergistic) effects of both drugs on the tumour cells were modelled through a nonlinear pharmacodynamics effects function *E*(*C*_*PAC1*_(*t*), *C*_*TRAIL*_(*t*)). This combined effect is given by

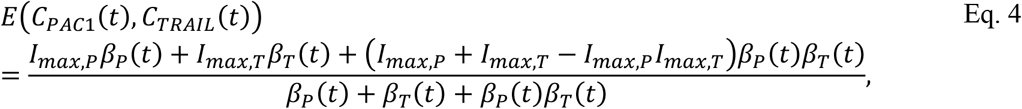

Where

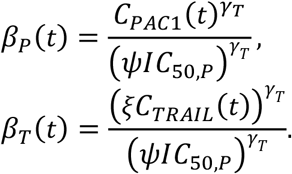

This model (**Eq. 4**) was originally conceived to describe the pharmacodynamic interactions *in vitro* of interleukin-10 and prednisolone^26^. In this model, *I*_*max, P*_ and *I*_*max, T*_ denote the maximum inhibitory effects achieved by PAC-1 and TRAIL, respectively, *IC*_50, *P*_ and *IC*_50, *T*_ are the half-effect concentrations of either drug (based on the classical Hill equation^27^), and *γ*_*P*_ and *γ*_*T*_ are the Hill coefficients.

The ratio of potency of each drug is expressed by *ξ* = *IC*_50, *P*_/*IC*_50, *T*_. In the case when one drug is zero, the *E*(*C*_*PAC1*_(*t*), *C*_*TRAIL*_(*t*)) simplifies to the well-known Hill function (or Michaelis-Menten equation when the Hill coefficient is equal to 1)^27^. A representative simulation of the effect function *E*(*C*_*PAC1*_(*t*), *C*_*TRAIL*_(*t*)) dependent on varying concentrations of PAC-1 and TRAIL is given in **Figure 1**B highlighting the previously-determined synergistic interaction.

The pharmacokinetic models in **Eqs. 2** and **3** can be rewritten to express drug elimination rates (*k*_*e*_, in units of per time) in terms of more conventional PK parameters, namely volume of distribution (*Vd*, in units of volume) and clearance (*CL*, in units of volume/time) using the conversion

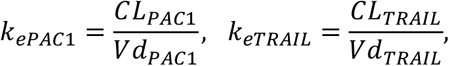

where *CL*_*PAC1*_ is the clearance and *Vd*_*PAC1*_ the volume of distribution for PAC-1, and *CL*_*TRAIL*_ and *Vd*_*TRAIL*_ are the clearance and volume distribution for TRAIL. Following previous animal and human PK studies^8,29,30^, we took PAC-1 and TRAIL doses to be 75mg and 3mg/kg, respectively.

### Parameter estimation

We used our previous in vitro experimental results3 to estimate average parameter values for the model in Eqs. **1** − **3**, and to define the values in the drug effects function (Eq. 4). We fit the tumour growth rate (*r*) and tumour carrying capacity (*K*) in the Gompertzian tumour growth term in Eq. 1 to the adult granulosa cell tumour KGN cell line cell counts. Estimated parameter values from Crosley et al.3 were used to describe the pharmacodynamics model. Model parameter values are given in **Table 1** and the resulting fit in **Figure S1** Supplementary Information.

**Table 1.** Model parameters. Details about parameter estimation are provided in the Methods.

| Parameter | Units | Description | Value | Reference |
| --- | --- | --- | --- | --- |
| $r$ | 1/day | Tumour cell proliferation rate | 0.052696 | Figure S1 |
| $K$ | | Tumour carrying capacity | $1.745 \times 10^9$ | Figure S1 |
| $C_0$ | cells | Initial number of tumour cells | 5000 | Crosley *et al.*<sup>3</sup> |
| $I_{max,P}$ | Dimensionless | Maximum PAC-1 efficacy | 1.0304* | Crosley *et al.*<sup>3</sup> |
| $I_{max,T}$ | Dimensionless | Maximum TRAIL efficacy | 0.5278 | Crosley *et al.*<sup>3</sup> |
| $IC_{50,P}$ | $\mu\text{M}$ | PAC-1 half-effect concentration | 40.3135 | Crosley *et al.*<sup>3</sup> |
| $IC_{50,T}$ | $\text{ng/mL}$ | TRAIL half-effect concentration | 15.1509 | Crosley *et al.*<sup>3</sup> |
| $\gamma_P$ | Dimensionless | PAC-1 Hill coefficient | 0.6829 | Crosley *et al.*<sup>3</sup> |
| $\gamma_T$ | Dimensionless | TRAIL Hill coefficient | 0.027865 | Crosley *et al.*<sup>3</sup> |
| $\psi$ | | Interaction strength for drugs | 0.8 | Crosley *et al.*<sup>3</sup> |
| $n_p$ | Patients | Number of virtual patients | 400 | -- |

### Implementation of an *in silico* clinical trial to determine combination dosing strategies

To reflect pharmacokinetic interindividual variability and the heterogeneity in treatment outcomes, we generated a virtual clinical trial using parameter estimates for patients sampled from real patient data (**Figure 2**A). Population data for both height and body mass index (BMI) for men and women in 200 countries are available from two recent surveys carried out by the NCD Risk Factor Collaboration^31,32^ and published online at http://ncdrisc.org/data-downloads.html. We used data for BMI and heights from women in the United States to generate individual virtual patient BMIs and heights by randomly sampling from normal distributions defined by reported mean and standard deviations. These values were then converted to body weight (BW) using the standard BMI formula

**Figure 2.**
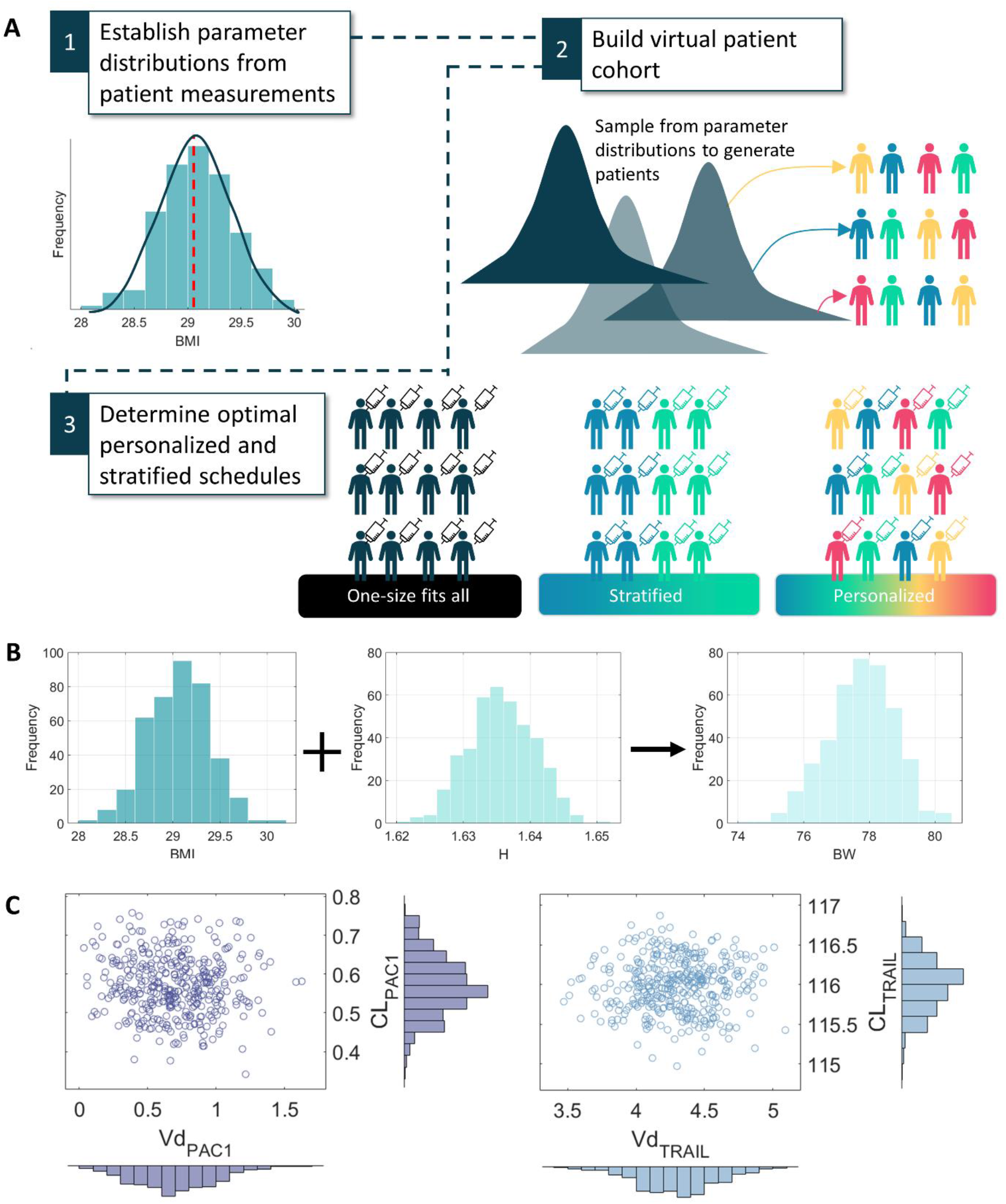
Generation and pharmacological characteristics of the 400 virtual patients. (A) Schematic describing the virtual clinical trial strategy. 1. Parameter distributions were established from measurements in the literature. 2. Virtual patient parameters were then sampled from parameter distributions established in Step 1. 3. Treatment protocols were optimized considering treatment individualization, stratified protocols determined by grouping subgroups with similar pharmacological characteristics, and a one-size fits all protocol. (B) Virtual patients body mass index, BMI, and height, H, distributions were used to determine body weight, BW. The distributions for these virtual patient characteristics are given in the denoted histograms. (C) Resulting distribution of pharmacokinetic parameters, *CL* and *Vd*, of PAC-1 (left) and TRAIL (right) for the virtual cohort. Marginal distributions for each parameter are plotted as histograms with individual samples of the bivariate distributions plotted as a scatter plot.

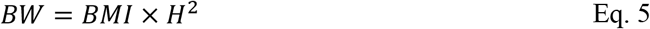

We sampled *n*_*p*_ = 400 patients using this scheme (distributions of BMIs, heights, and body weights is provided in **Figure 2**B). Body weight was used to determine the total dose size of TRAIL.

Using existing population PK data, we also determined patient-specific clearance rates and volumes of distribution for each drug. For PAC-1, the average total body clearance and central volume of distribution (*Vd*) was previously reported to be 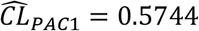 with variance *σ*_*PAC1*_ = 0.0757, and 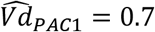 with variance *σ*_*PAC1*_ = 0.3 ^8^, respectively. Individual patient clearance *CL* and *Vd* for PAC-1 were then sampled from a multivariate normal distribution of these two parameters (**Figure 2**C). Similarly, the total body clearance and volume of distribution (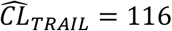 and variance *σ*_*TRAIL*_ = 0.315, and 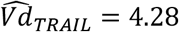 and *σ*_TRAIL_ = 0.301^33^) were used to define individual patient clearance and volume distribution values for TRAIL (**Figure 2**C). All parameter samples were restricted to being non-negative.

For our cohort of 400 virtual patients, we then individualized combination PAC-1 and TRAIL regimens in an *in silico* clinical trial, as in previous work^16,21,34–37^. For this, we assumed that both drugs were given on one-week cycles and restricted each dose size to be a multiple of the two standard doses we considered (75mg for PAC-1 and 3mg/kg for TRAIL). For each patient, we then used a genetic algorithm^21,22,38^ to determine whether or not a PAC-1/TRAIL dose was given on each possible administration day (once a week) and, if given, the appropriate dose size (up to 10 times the standard dose, within previously investigated dosing ranges). We considered treatment regimens over a total of 10 weeks. Extending this analysis, we then applied a genetic algorithm to determine a stratified cohort protocol (i.e., a cohort for patients in specific subgroups, see Results) and a one-size-fits-all protocol (**Figure 2**A).

### Numerical simulations

All simulations were run using Matlab 2020b^39^. The function *ode45* was used to solve the model. Parameter values were fitted using *lsqnonlin* (non-linear least squares optimization), and the function *ga*^40^ was used to run the genetic algorithm (**Figure S2** Supplementary Information) to optimize treatment schedules. A kernel density estimate (KDE) was used to estimate the distribution for the subgroups within our cohort of virtual patients using Matlab’s *ksdensity*. Matlab’s *random* was used to draw from the KDE for the probability distribution function (PDF) of each parameter. To approximate tumour eradication, we assumed complete cell elimination occurs when the number of cells *C*(*t*) is less than 1 (i.e. *C*(*t*) < 1).

To determine optimal dosage regimens, we considered PAC-1 and TRAIL doses to be given every week on the first day of the week from week 0 through to week 10. In every week *i*, the size of the PAC-1 and TRAIL dosages were determined by the *i*th entry of the vectors ***Z***_*P*_ and ***Z***_*T*_, i.e. *Z*_*P, i*_ and *Z*_*T, i*_, where ***Z***_*P*_ and ***Z***_*T*_ represents the dose size multiple for week *i* (ranging from 0 to 10). In other words, *Z*_*P*, 3_ is the PAC-1 dose multiple for week 3. We then minimized the objective function:

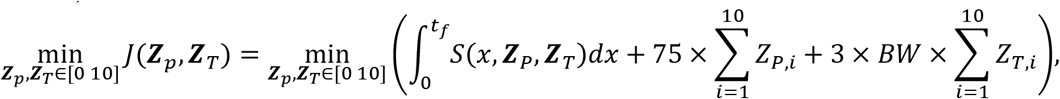

where the first term represents the total tumour burden, and the second and third terms the total amount of PAC-1 and TRAIL administered, respectively. In this way, each virtual patient’s regimen was optimized to simultaneously reduce their tumour size using the minimal amount of each drug. Note that *Z*_*P, i*_ and *Z*_*T, i*_ are integers between 0 and 10.

To validate our optimal protocols suggested for subgroups, we used KDE to generate a new cohort of patients with equal numbers of patients from each subgroup. Three optimal protocols were then determined: the full cohort optimal (one-size-fits-all), the subgroup optimal protocol (stratified) and the individual patient optimal (personalised), see **Fig 2**A. The full cohort optimal protocol was determined by applying a genetic algorithm to minimise the objective function above for all patients simultaneously. Similarly, the subgroup optimal protocol was determined by minimising the objective function for all patients in a subgroup. Finally, the individualised protocol was determined by optimising the objective function for each virtual patient individually.

## Results

### Variability in tumour growth dynamics are driven primarily by PAC-1 pharmacokinetics

To obtain average estimates for the growth rate (*r*) and carrying capacity (*K*) of KGN cells (**Table 1**), we fit a Gompertzian tumour growth model (**Eq. 1**) to KGN cell counts (**Figure S1**). Virtual patients (n=400) were then created by sampling from distributions (**Figure 2**A) centred around the estimated parameters for ***BW***, *Vd*_*PAC1*_, *CL*_*PAC1*_, *Vd*_*TRAIL*_ and *CL*_*TRAIL*_ (**Figure 2**B-C), as described in the Methods. The resulting distributions for the initial PAC-1 dose (*C*_*PAC1*_(0)) and PAC-1 and TRAIL elimination rates, *k*_*ePAC1*_ and *k*_*eTRAIL*_, were generated using each patients individual ***BW***, *Vd*_*PAC1*_, *CL*_*PAC1*_, *Vd*_*TRAIL*_ and *CL*_*TRAIL*_ value (**Figure 3**A-C).

**Figure 3.**
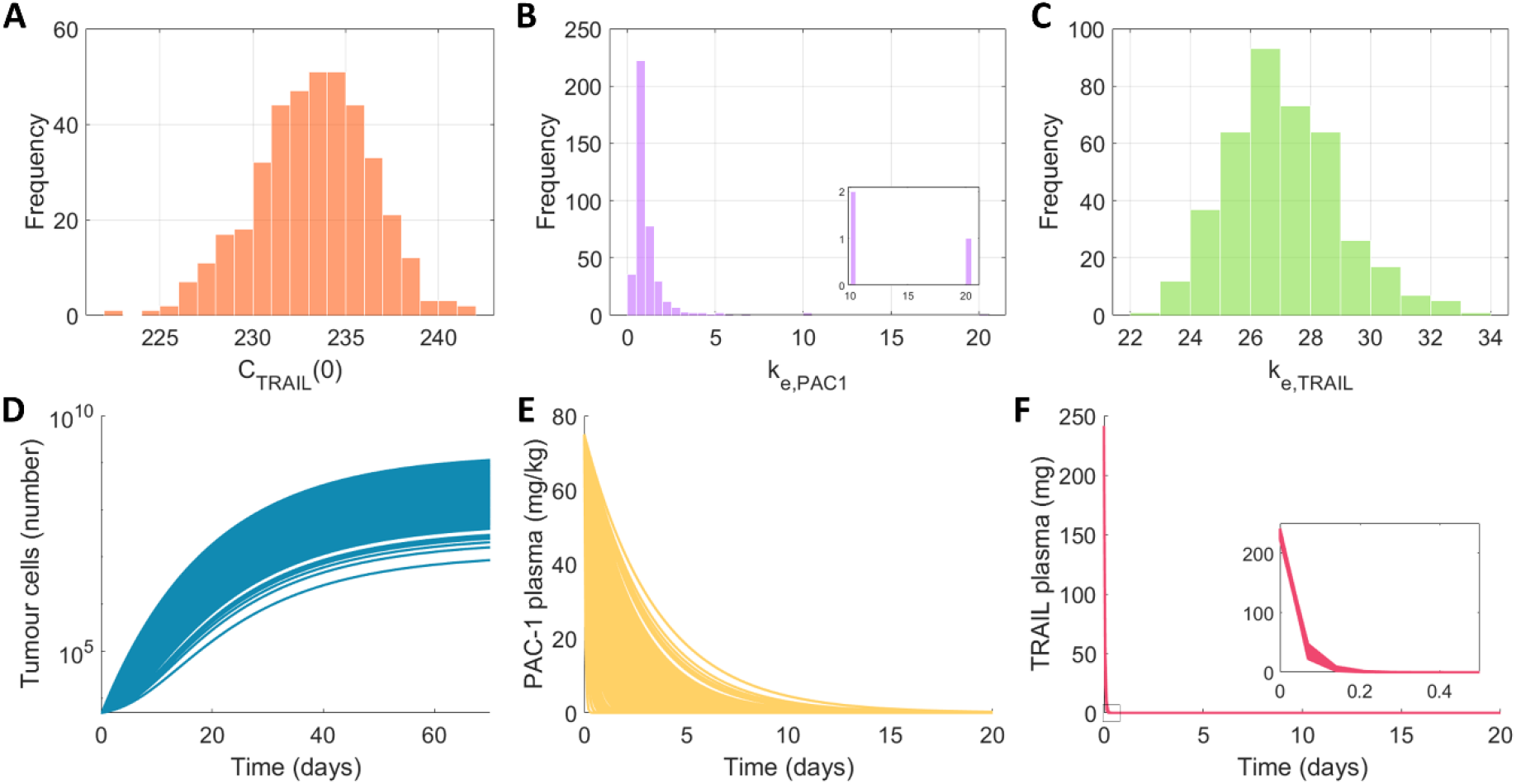
Baseline distributions of pharmacokinetic and pharmacodynamic responses in the virtual cohort. (A-C) Histograms of the distributions key pharmacokinetic parameters in the virtual cohort. (A) The distribution for initial TRAIL dose (*C*_*TRAIL*_(0)) determined based on each virtual patient’s BW. (B) Distribution of PAC-1 elimination rates (*k*_*ePAC1*_). (C) Distribution of TRAIL elimination rates (*k*_*eTRAIL*_). (D-F) Responses in the virtual cohort after a single dose of PAC-1 (*C*_*PAC1*_(0) = 75) and TRAIL (*C*_*TRAIL*_(0) = 3 × *BW*). For the 400 virtual patients the output of the model is plotted for (D) the number of cancer cells (*G*(*t*)), (E) the concentration of PAC-1 (*C*_*PAC1*_(*t*)), and (F) the concentration of TRAIL (*C*_*TRAIL*_(*t*)).

To assess baseline plasma concentrations following PAC-1 and TRAIL administrations along with tumour cell counts, we simulated a single IV bolus of both drugs in the virtual patient cohort (**Figure 3**D-F). The variability in drug concentrations produced large variations in the dynamics of tumour cell growth (**Figure 3**D). We also found that patient-specific PK parameter differences resulted in large variations in PAC-1 plasma concentrations (**Figure 3**E) and only limited variations in TRAIL concentrations (**Figure 3**F), which can be potentially attributed to the rapid elimination rate of TRAIL compared to PAC-1 (**Figure 3**B-C).

### Individualized combination PAC-1 and TRAIL schedules predicted to be highly effective in a portion of the virtual patient cohort

The effect of combination PAC-1 and TRAIL therapy in patients is unknown, and viable treatment schedules remain to be determined. Based on our previous *in vitro* study^3^, we considered weekly administrations of both drugs for 10 weeks (**Figure 4**A) to aid in the preclinical translation of combination PAC-1 and TRAIL for GCT. Accordingly, at the beginning of the *i*th week, we allowed for a PAC-1 dose of size 75 × *Z*_*P, i*_ and a TRAIL dose of size 3 × *BW* × *Z*_*T, i*_ to be administered (note that either or both *Z*_*P, i*_ and *Z*_*T, i*_ could also be equal to 0). Each individual patient’s optimal treatment schedule was then determined using a genetic algorithm, as described in the Methods.

**Figure 4.**
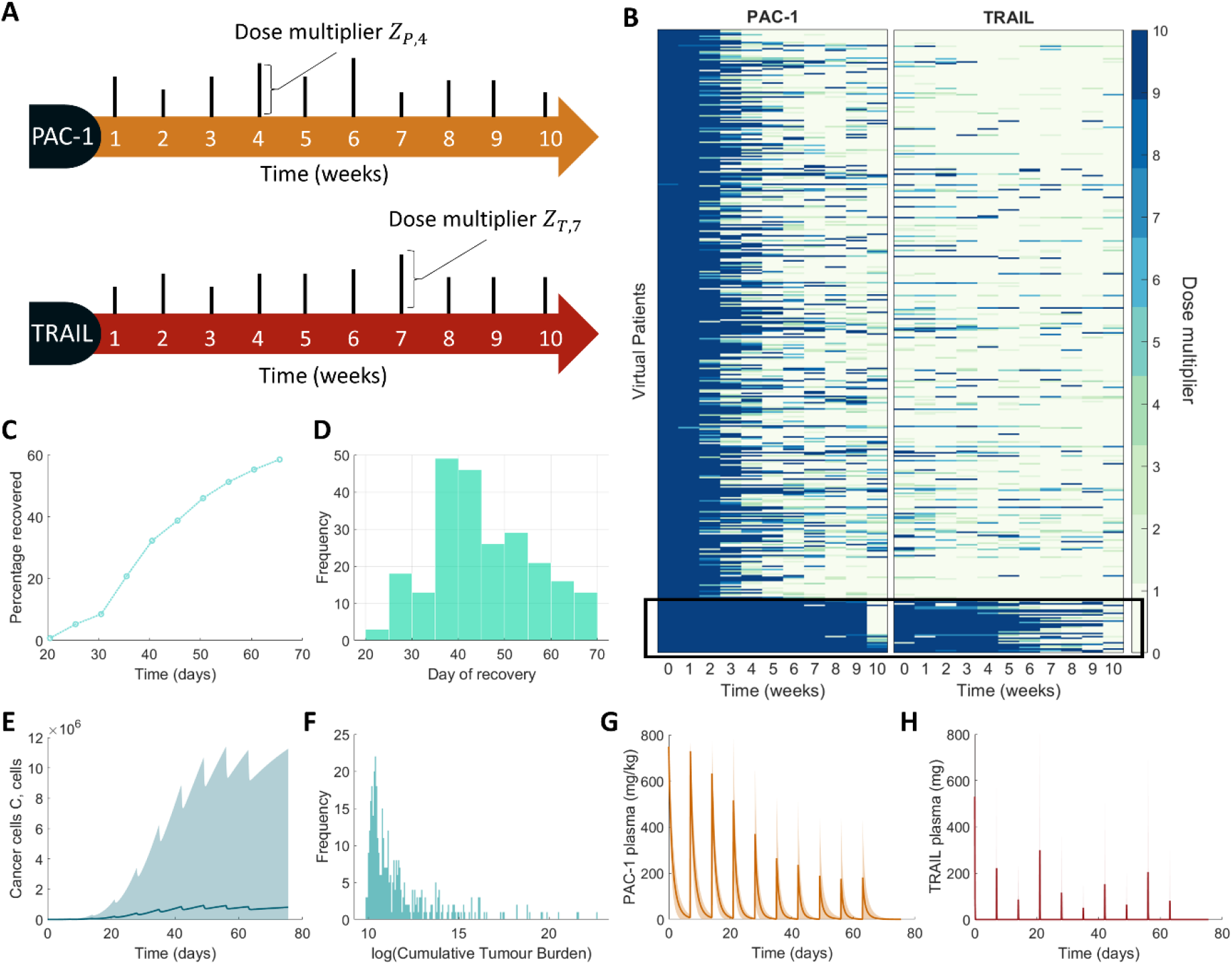
Optimal individualized combination PAC-1 and TRAIL treatment schedules. (A) PAC-1 and TRAIL were administered simultaneously every week for 10 weeks at a weekly dosage multiplier of *Z*_*P, i*_ and *Z*_*T, i*_ for week *i*, respectively, from the base dosage. Weekly dose sizes were allowed to vary week to week for each patient. (B) Optimal individual regimens for each virtual patient. Dose size is given as a multiple of the standard dosing of either drug in shades of green to blue, where light green is no dose and dark blue is a 10-fold increase. The *i*th column corresponds to the dosage size on the *i*th week and the *k*th row corresponds to the *k*th patient’s optimal protocol. Patients have been ordered to highlight those requiring the highest dosages of PAC-1 (solid black box). (C)-(D) Percentage of the population that has recovered cancer cell elimination) and the day of recovery is given. (E)-(H) Model dynamics for the cohort under their optimal protocol. (E), (G)-(H) Solid line and shaded regions representing the mean and standard deviation in the full cohort. (F) Histogram of log cumulative tumour burdens.

Using our regimen optimization scheme, we found that most virtual patients required large doses of PAC-1 early in the regimen (weeks 1 and 2) whereas there was no clear pattern to the optimal TRAIL dosage schedule across the cohort (**Figure 4**B). Ordering the patients by total PAC-1 dosage, however, we noticed a clear pattern: patients who required highest dosages of PAC-1 also required higher dosages of TRAIL (**Figure 4**B). We further predicted that the majority of patients (≈ 60%) could achieve remission (cancer cell elimination) over the treatment period under their personalized regimen (**Figure 4**C), with most patients recovering early. However, we found significant variations in the time of recovery (**Figure 4**D). The number of tumour cells remaining for some patients was quite high (**Figure 4E**), with the cumulative tumour burden reaching extremely high values for some patients (**Figure 4**F). This suggests that this combination therapy is ineffective for some patients and other treatment regimens are necessary. PK dynamics for PAC-1 and TRAIL (**Figure 4**G-H) support the idea that large PAC-1 doses are optimal for most patients in the cohort, whereas TRAIL dosages show no consistent pattern.

### Optimal individualized treatment schedules are determined by PAC-1 pharmacokinetic parameters

Based on the individualization results, we next sought to determine whether we could distinguish specific patient subgroups predicted to require different treatment regimens or establish characteristics of responders/nonresponders^16,21^. By systematically analysing the results in **Figure 4B**, we found a subgroup of patients requiring higher dosages of PAC-1 and TRAIL (**Figure 5**A) who exhibited poor outcomes with respect to tumour growth (**Figure 5**B-C) compared to the rest of the cohort. Further examination revealed that patients requiring higher dosages of PAC-1 also had higher PAC-1 elimination rates (**Figure 5**D-F), so we called this subgroup the High PAC-1 elimination subgroup. Other key PK parameters, including BWs and TRAIL elimination, remained consistent between each subgroup (**Figure 5**E and F for the individual distributions of the clearance and volume distribution that were used to calculate these clearance rates, and **Figure S3**A-D in the Supplementary Information for further subgroup dynamics. Dynamics for the other model variables for each group can also be found in **Figure S3**E-F in the Supplementary Information). We therefore divided our virtual cohort into two subgroups of patients: High PAC-1 elimination cohort and Normal PAC-1 elimination cohort.

**Figure 5.**
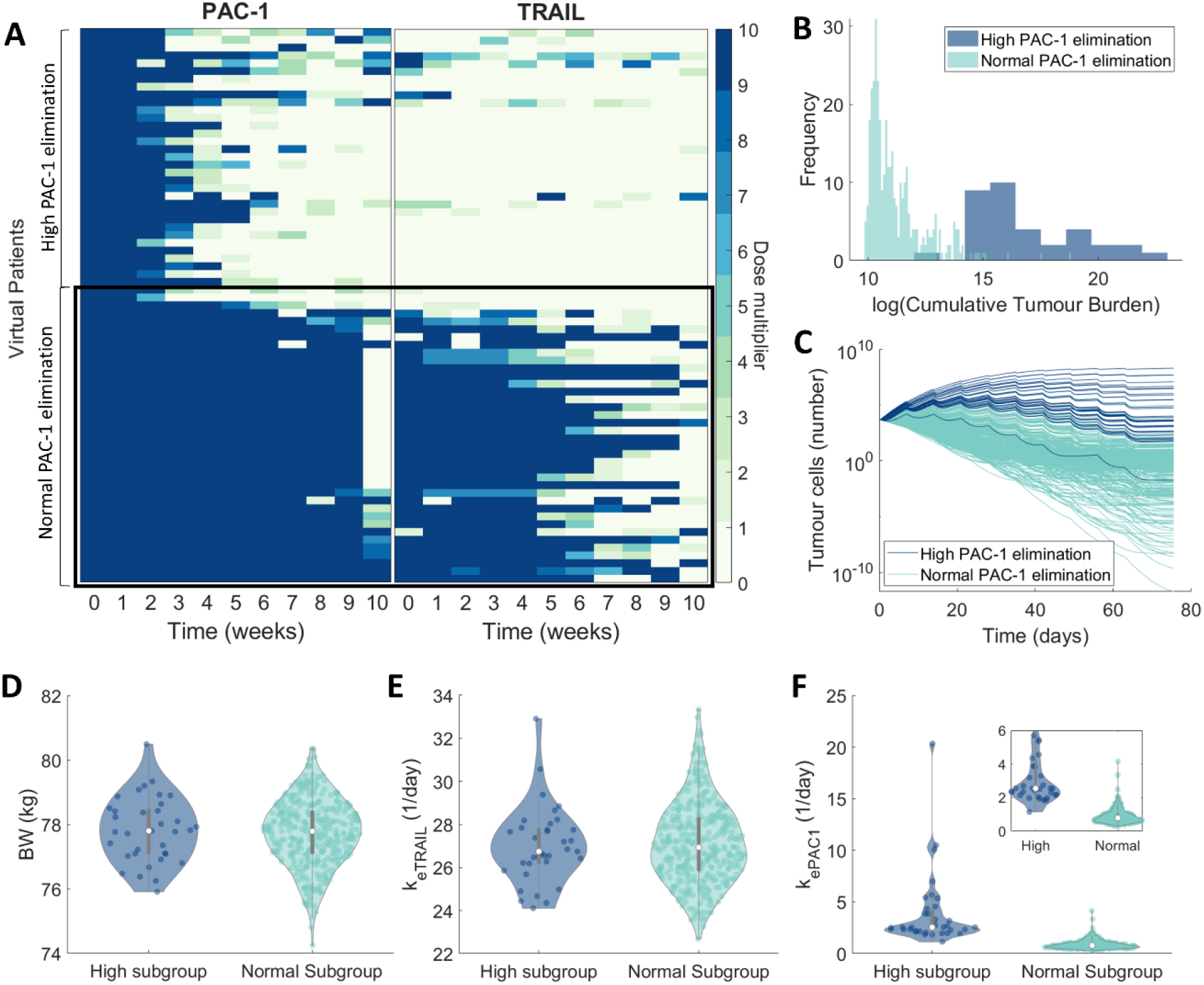
PAC-1 clearance rates distinguish poor responders in virtual patient cohort. (A) Zooming in on the bottom 100 patients’ optimal individual protocols in **Figure 4**B, the difference between individualized protocols distinguishes the High PAC-1 and Normal PAC-1 elimination subgroups. (B) Histogram of the cumulative tumour burdens and (C) trajectories for individual patient tumour cell counts in both subgroups. Identifying characteristics of virtual patients in the High and Normal PAC-1 elimination rate subgroups according to (D) body weight (*BW*), (E) TRAIL elimination rate (*k*_*eTRAIL*_), and (F) PAC-1 elimination rate (*k*_*ePAC1*_). The violin plots for the specific clearance and volume distribution for both subgroups can be found in **Figure S3**A-D in the Supplementary Information.

Reordering the individualized protocols for weekly doses of PAC-1 and TRAIL (**Figure 4**B) with respect to both the High PAC-1 and Normal PAC-1 elimination subgroups highlights clear differences between protocols in each group (**Figure 5**A). Notably, virtual patients in the High PAC-1 elimination subgroup required much higher doses of both PAC-1 and TRAIL in their optimal protocols than virtual patients with “normal” PAC-1 elimination. Further, examining the dynamics of each cohort in terms of the cumulative tumour burden (**Figure 5**B) and tumour cell counts over time (**Figure 5**C), our model predicted that patients in the Normal PAC-1 elimination subgroup achieved, on average, a lower tumour size under their optimal protocols than those in the High PAC-1 elimination subgroup. This suggests that the Normal PAC-1 elimination subgroup is comprised of virtual patients who responded more robustly to treatment. Further, there were no patients in the High PAC-1 elimination subgroup who achieved complete clearance of their tumours.

### High PAC-1 elimination rates translate to low therapeutic efficacy

While individualized protocols are an exciting area of cancer treatment, feasibility is a barrier to clinical implementation. Given the differences between optimal individualized protocols for virtual patients in the High vs. Normal PAC-1 elimination subgroups (**Figure 5**A), we tested whether robust optimal treatment protocols existed for patients after segregation based on PAC-1 clearance pharmacokinetics. For this, we generated two new cohorts of 200 virtual patients each based on the parameter distributions identified for the High and Normal PAC-1 elimination subgroups (**Figure 5**D-F and **Figure S3**A-D Supplementary Information). For each new virtual patient, we sampled from a kernel density estimate for the probability density function for *Vd*_*PAC1*_, *CL*_*PAC1*_, *Vd*_*TRAIL*_, *CL*_*TRAIL*_, and *BW*, based on the parameters of individuals in the subgroups (**Figure 6**A). We verified that parameter values for these newly generated virtual patients were comparable with those in both the High and Normal PAC-1 elimination subgroups using a Kolmogorov-Smirnov test at the 5% significance level (**Figure S4**A Supplementary Information). We then plotted the characteristics of the subgroups to confirm they matched the original subgroups (**Figure S4**B Supplementary Information), i.e. the new High PAC-1 elimination cohort had low *Vd*_*PAC1*_ compared to the new Normal PAC-1 elimination cohort.

**Figure 6.**
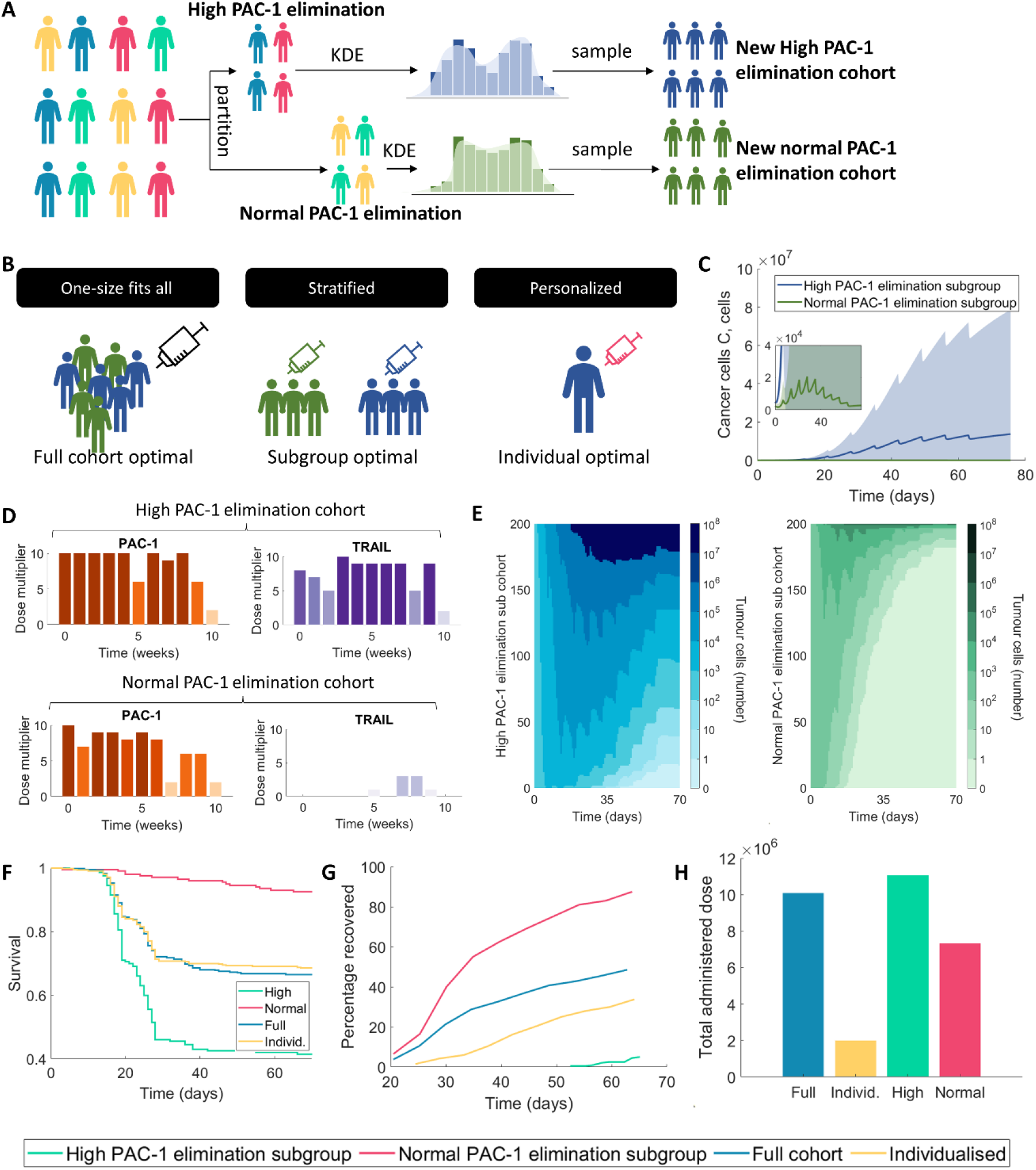
Characteristics of responders to combination PAC-1 and TRAIL therapy are driven by PAC-1 elimination rates. (A) Two new virtual cohorts were generated using Kernel Density estimates for the probability distribution functions for parameters for the High PAC-1 elimination cohort and the Normal PAC-1 elimination cohort (**Figure S4** Supplementary Information). (B) Schematic describing the three optimal protocols determined for the new cohort: full cohort optimal (one-size-fits-all), subgroup optimal (stratified) and individual optimal (personalised), see **Fig 2**A. (C-E) Optimal protocols for the High PAC-1 elimination cohort and Normal PAC-1 elimination were determined using our *in silico* clinical trial approach (see Methods). (C) Average and standard deviation of the number of tumour cells of each subgroup under the subgroup optimal protocols. (D) Weekly optimal dose multipliers for both PAC-1 and TRAIL for the High PAC-1 elimination subgroup (top) and the Normal PAC-1 elimination subgroup (bottom) protocol. (E) Individual patient trajectories under subgroup optimal protocols. Each virtual patient’s total number of tumour cells over 70 days is visualized as a heatmap, where the *i*th column corresponds to the *i*th day and the *j*th column corresponding to the *j*th patient. (F-H) Comparison of the treatment efficacy for the new virtual cohort under four different protocols: full cohort level optimized protocol (**Figure S6** Supplementary Information), individualized schedules (**Figure S5** Supplementary Information), subgroup-optimal protocols, (i.e. High PAC-1 and Normal PAC-1 elimination subgroups). (F) Comparison of Kaplan-Meier survival under the different protocols considered. (H) Recovery percentage (Tumour cells <1) over the course of treatment under the four different protocols. (G) Total dose burden for all patients under the four protocols.

To thoroughly investigate the possible treatment options for High PAC-1 elimination and Normal PAC-1 elimination patients, we then determined three different possible treatment schedules (**Figure 6**B): full cohort optimal (one-size-fits-all), subgroup optimal (stratified), individual optimal (personalised). The full cohort optimal schedule was a single treatment schedule optimal for all patients in the new cohort simultaneously using a genetic algorithm. Similarly, the subgroup optimal schedule was a single treatment schedule optimal for all patients in one subgroup, where the two subgroups might have different optimal schedules. For the individual schedule, as before, we then individualized treatment schedules for patients in the new subgroups using a genetic algorithm under the same dosing constraints (**Figure S5** Supplementary Information). We again found that both PAC-1 and TRAIL doses in the High PAC-1 elimination group were higher than those in the Normal PAC-1 elimination subgroup, impacting slightly on average tumour growth dynamics in both groups (not shown).

After confirming the similarity of each subgroup’s characteristics to the original cohort, we then sought to determine whether we could find a single optimal protocol for all patients in the High PAC-1 elimination cohort and single optimal protocol for all patients in the Normal PAC-1 elimination cohort: the subgroup optimal (**Figure 6**C-E). Using our same *in silico* clinical trial approach, we determined a stratified protocol for each cohort (**Figure 6**D). Under these protocols, we found that patients in the Normal PAC-1 elimination cohort had lower on average tumour volumes than those in the High PAC-1 elimination cohort (**Figure 6**C), as before. Furthermore, examining the individual tumour cell count for each patient under this new protocol revealed most patients in the Normal PAC-1 elimination subgroup reached a state of cancer cell elimination (**Figure 6**E). In comparison, patients in the High PAC-1 elimination subgroup had much larger tumour cell counts on day 70 after treatment with their subgroup’s optimal protocol.

Given a number of patients were responsive to the subgroup optimal protocol, we next sought to determine a full cohort optimal (one-size-fits-all) treatment (**Figure S6** Supplementary Information). To evaluate the effectiveness of this protocol against the other subgroup protocols and individualised protocol, we compared predicted virtual patient tumour evolution under the three different treatment protocols (**Figure S7** Supplementary Information). We found the subgroup protocol for Normal PAC-1 elimination rates showed the most substantial reduction in tumour growth.

We next compared predicted virtual patient survival under their optimized individual protocol, the relevant subgroup protocol, and the optimal schedule for the combined cohort. To measure survival in our virtual patient cohort and compare different schedules, we introduced a proxy based on clinical evidence that tumour size is a highly prognostic factor for reduced survival probability in ovarian granulosa tumours over 10cm^41,42^. Previous models for melanoma and breast cancer have assumed ~10^6^ cells/mm^3 43,44^, assuming ovarian cancers are less dense in comparison we assumed there are ~10^3^ cells/mm^3^ and that a tumour with radius >5cm is deadly^41,42^. With this, we constructed a probability density function for patient death using a shifted exponential, where survival probability is 1 for a tumour with 0 cells and 0 for a tumour close to 10^8^ cells (**Figure S8** Supplementary Information). We found that patient survival for the High PAC-1 elimination subgroup under the corresponding stratified protocol had the worst survival (**Figure 6**F). Interestingly, both the Full cohort optimal protocol and the individualized protocols had qualitatively similar survival projections, suggesting there was no benefit to treating with an individualized schedule over a one-size-fits-all protocol. The best survival was seen with the Normal PAC-1 elimination cohort treated with its corresponding stratified protocol.

Quantifying the percentage of the population that had recovered under the different protocols (**Figure 6**G), we found that almost all patients in the Normal PAC-1 elimination cohort treated with their cohort’s protocol recovered. Interestingly, the next most effective protocol was seen in the full cohort protocol, where ≈ 60% of patients recovered. The reduction in efficacy of the individualized protocol for this new cohort compared to the original cohort (**Figure 4**C) is due to the higher proportion of patients in the High PAC-1 elimination cohort for which treatment is less effective.

Further, the total drug burden in each of the three protocols was virtually unchanged (**Figure** 6H), suggesting that stratifying patients by PAC-1 elimination and then only treating Normal PAC-1 elimination patients with a cohort protocol may be the most effective way to provide efficacy with regards to survival using combination PAC-1 and TRAIL regimens.

## Discussion

Designing and developing novel approaches to cancer treatment is a lengthy and costly process with high rates of attrition^45,46^. The need for new approaches to treating ovarian cancer, including rare subtypes like granulosa cell tumour of the ovary, is highlighted by the lack of robust treatment options for advanced and late-stage disease^2^. We have previously established that PAC-1 and TRAIL, both apoptosis-activating molecules, act synergistically *in vitro* to induce KGN cell death by combining a mathematical model and experimental results^3^. In this paper, we expanded on our model to run *in silico* clinical trials of combination PAC-1 and TRAIL with the aim of establishing the viability of this combined therapy in GCT.

After integrating simple pharmacokinetic models for both PAC-1 and TRAIL to our synergistic effects model, we calibrated our mathematical model to experimental data of KGN cell growth *in vitro*. We then leveraged the model to expand a virtual patient cohort according to PK variability data. Individualization of combination PAC-1 and TRAIL treatment schedules identified a distinct subgroup within the cohort who responded particularly poorly to treatment and had markedly different individualized regimens than the rest of the virtual patients. In this High PAC-1 elimination subgroup, PAC-1 clearance rates were much higher than those in the Normal PAC-1 elimination group, suggesting that PAC-1 pharmacokinetics may be an exclusionary factor for combination PAC-1 and TRAIL therapeutic approaches. The idea of stratifying human patients for PAC-1 was previously discussed in the context of central nervous system cancers^47^.

To validate this prediction, we generated a new virtual patient cohort with an even distribution of patients in the High PAC-1 and Normal PAC-1 elimination subgroups. In the new cohort, we investigated three regimen optimization strategies: individualization (as before), stratification, and one-size-fits-all. In all three cases, the High PAC-1 elimination subgroup performed worse than the Normal subgroup, validating our findings in the original cohort. Overall, patients in the Normal PAC-1 elimination subgroup treated with their specific (stratified) treatment regimen performed well under treatment and did not require individualization for therapeutic efficacy. However, of the four optimization scenarios (individualization, one-size-fits-all, and the two stratified regimens), we predicted the worst outcomes in the High PAC-1 elimination subgroup with their subgroup-specific regimen. Together, these results reinforce that PAC-1 pharmacokinetics are critically important to this combined regimen’s success in patients and suggests the exclusion of patients with high PAC-1 clearance rates.

There are limitations to our study. PAC-1 and TRAIL remain investigational cancer drugs, and further studies are needed to establish improved pharmacokinetic models in humans. As we were focused on pharmacokinetic variability, we assumed a fixed initial number of GCT tumour cells per patient. Future work should examine the impact of physiological variability (beyond body weight heterogeneity). Further, our model of tumour growth was calibrated to *in vitro* data which is not fully representative of growth *in vivo* owing to the difficulties of studying GCT in murine models, as mice either lack the FoxL2 mutation frequent in adult GCT^48^ or are not very supportive of engraftment. Future improvements to the model include a more detailed description of the tumour microenvironment (represented here more simply through Gompertzian growth).

Nonetheless, this work highlights the importance of mathematical and PK/PD modelling in preclinical oncology research. Our TRAIL and PAC-1 virtual patient trials suggest key characteristics of GCT patients for whom combination PAC-1 and TRAIL therapy could be a viable treatment strategy. The continued study of new therapies to treat GCT and other ovarian cancers using quantitative approaches is a critical component of efforts to improve treatment and provide patients with effective and durable therapies.

## Funding

OC was supported by the Canada Research Chair in Differential Geometry and Topology. MC was funded by a J1 Research Scholar Grant from the Fonds de recherche du Québec-Santé. ALJ and MC were supported by NSERC Discovery Grant RGPIN-2018-04546 and funds from the Fondation du CHU Sainte-Justine to MC.

## Code availability

Code for the simulations in this manuscript can be found at https://github.com/AdrianneJennerQUT/virtual-clinical-trial-PAC1-TRAIL.

## Notes

### Competing Interest Statement

The authors have declared no competing interest.

