## Supplementary Information for "Establishing combination PAC-1 and TRAIL regimens for treating ovarian cancer based on patient-specific pharmacokinetic profiles using *in silico* clinical trials"

Contains:

- **Figures S1 – S8:** Supplementary figures for the results in the Main Text

### Supplementary Information

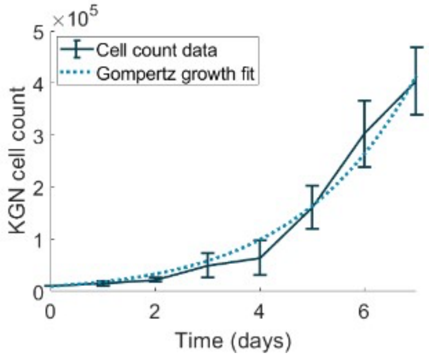


**Figure S1. Gompertzian tumour growth fit to KGN cell count measurements.** To determine the underlying tumour growth rate for the model, the Gompertz growth model was fit to data from an *in vitro* cell count assay for KGN cells (n=6). Parameter values from the fit can be found in **Table 1**.

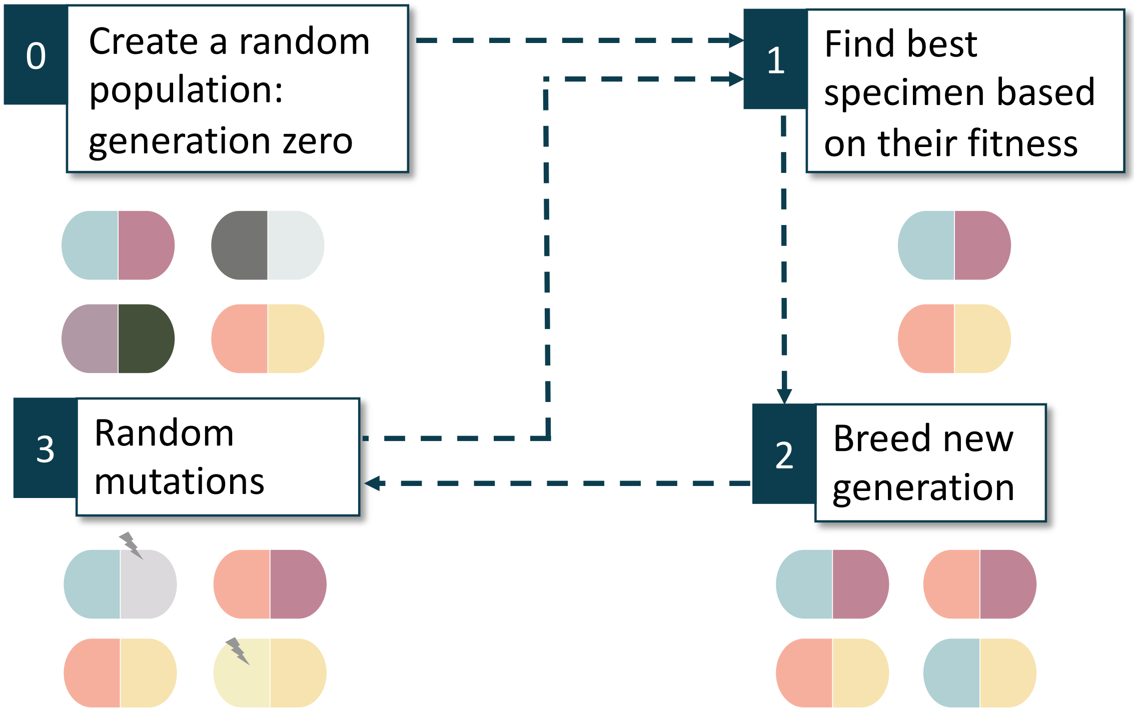


**Figure S2 Schematic depicting genetic algorithm used to optimize treatment protocols.** In general, the algorithm begins by generating a number of random possible treatment protocols, i.e. a population (or generation) of $\boldsymbol{Z_P}$ and $\boldsymbol{Z_T}$ vectors. The individuals in the generation are evaluated for their fitness. In other words, each treatment protocol in the generation is evaluated for the resulting tumour burden and total dose. In the case of the individualised protocol, this is only done for a single patient, whereas for the subgroup and full cohort optimal protocols, each treatment protocol is simulated for everyone in the subgroup/full cohort. The best possible protocols in the current generation are chosen and new generations are created from these. Some examples within this new generation are then randomly mutated and the process continues.

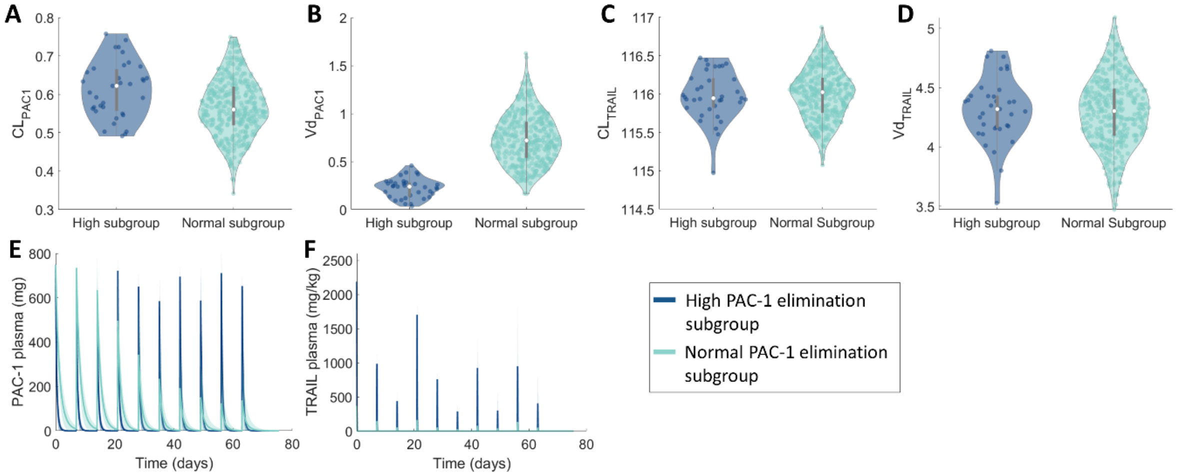


**Figure S3. Characteristics of subgroups of the virtual clinical patient population.** (A-D) Violin plots for the parameters $D_{PAC1}, CL_{PAC1}, VD_{TRAIL}, CL_{TRAIL},$ and $BW$ for patients in the High and Normal PAC-1 elimination subgroups for (A) $CL_{PAC1}$, (B) $Vd_{PAC1}$, (C) $CL_{TRAIL}$, and (D) $Vd_{TRAIL}$. Patient specific PAC-1 and TRAIL elimination rates ($k_{eTRAIL}$ and $k_{ePAC1}$, respectively) along with each patient's body weight are provided in **Figure 5Error! Reference source not found.**D-F. (E-F) Average model dynamics in the High and Normal PAC-1 elimination subgroups with respect to (E) PAC-1 concentrations, and (F) TRAIL concentrations under individualized regimens. Mean are indicated by the solid lines and standard deviation by shaded regions. The cancer cell dynamics are presented in **Figure 5**B-C.

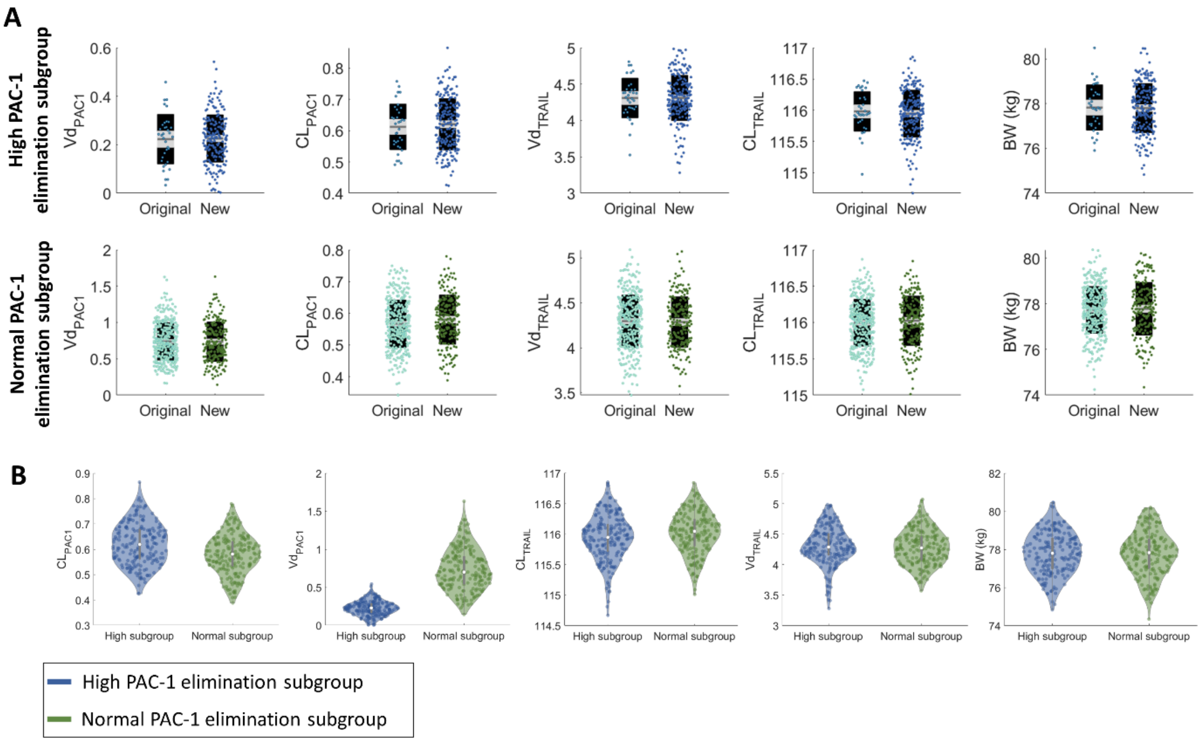


**Figure S4. New virtual cohort characteristics built from identified High and Normal PAC-1 elimination subgroups.** (A) Sampling the kernel density estimation to each subgroup's parameters (i.e. $Vd_{PAC1}$, $CL_{PAC1}$,$Vd_{TRAIL}$, $CL_{TRAIL}$, and $BW$, see **Figure 6**A), new virtual cohorts (n=200) were generated. The comparison between parameters for the High and Normal PAC-1 elimination subgroups and their corresponding newly sampled counterparts are given as box plots overlayed with a scatter plot of each virtual individual's parameters values. (B) Violin plots comparing each of the new subgroups' parameters.

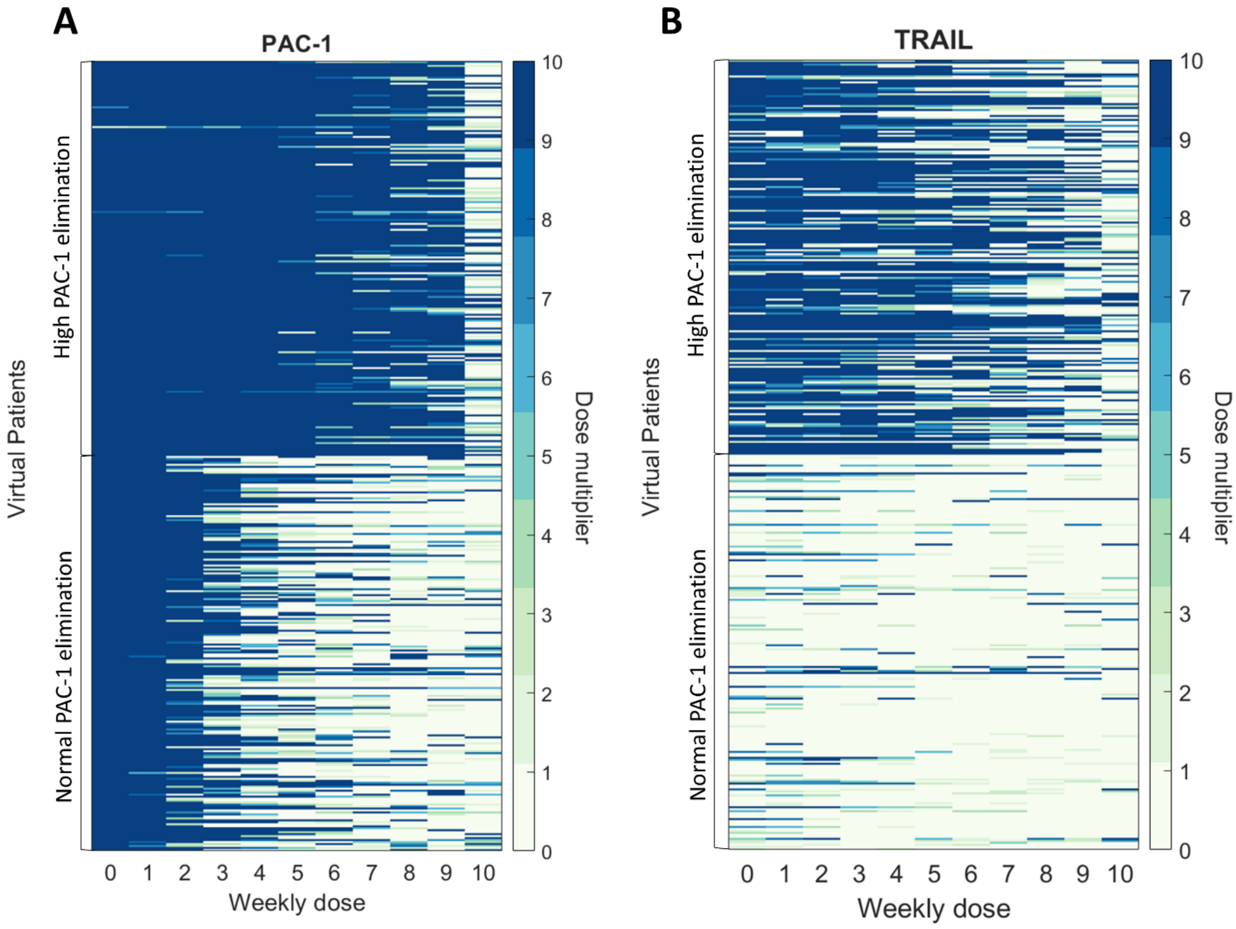


**Figure S5. Individualized protocol for new virtual patient cohort.** Individual regimens for each virtual patient were determined for (A) PAC-1 and (B) TRAIL, dosed every week for 10 weeks (see Main Text Methods and **Figure 4Error! Reference source not found.**A). Dose size is indicated as a multiple of the standard dosing of either drug. In both A and B, the $i$th column corresponds to the dose size on the $i$th week and the $k$th row corresponding to the $k$th patient's optimal protocol. The patients in the High PAC-1 elimination subgroup and Normal PAC-1 elimination subgroup have been noted.

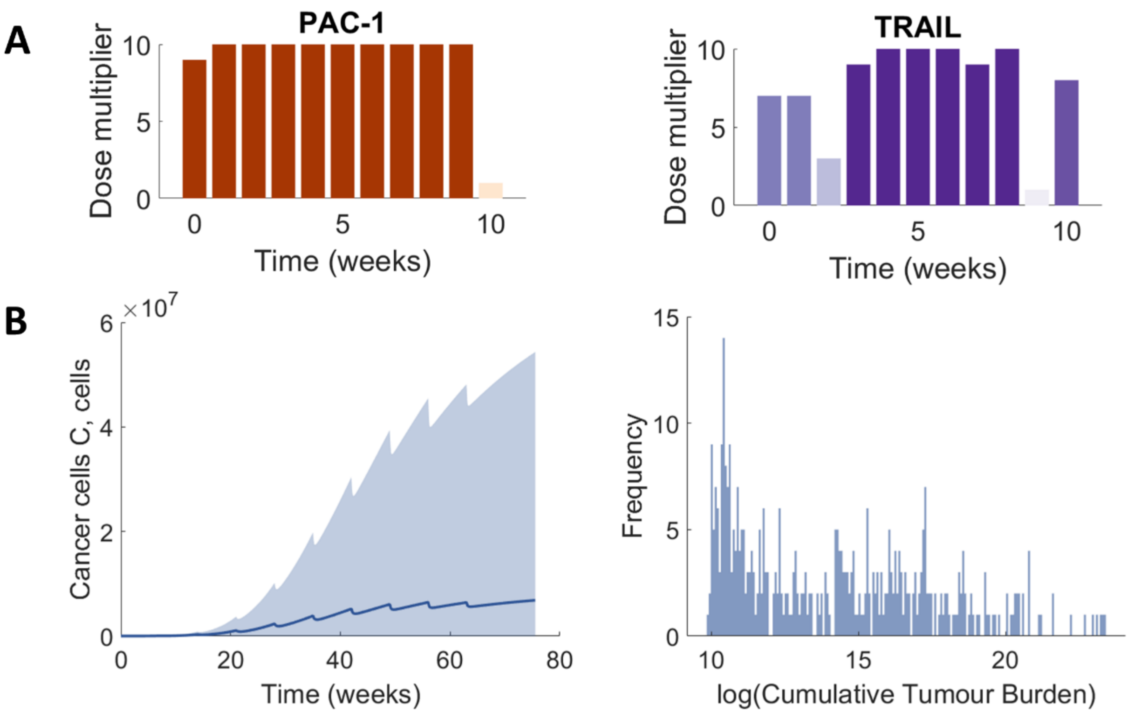


**Figure S6 Full cohort optimal PAC-1 and TRAIL dosage protocols.** (A) Optimal weekly dosage increase for PAC-1 and TRAIL for all virtual patients ($n = 400$) in the newly generated High PAC-1 elimination subgroup ($n = 200$) and the Normal PAC-1 elimination subgroup ($n = 200$). The colour of the bar corresponds to the increase on the dosage size. (B) The average and standard deviation for the cohort cancer cells and the cumulative tumour burden.

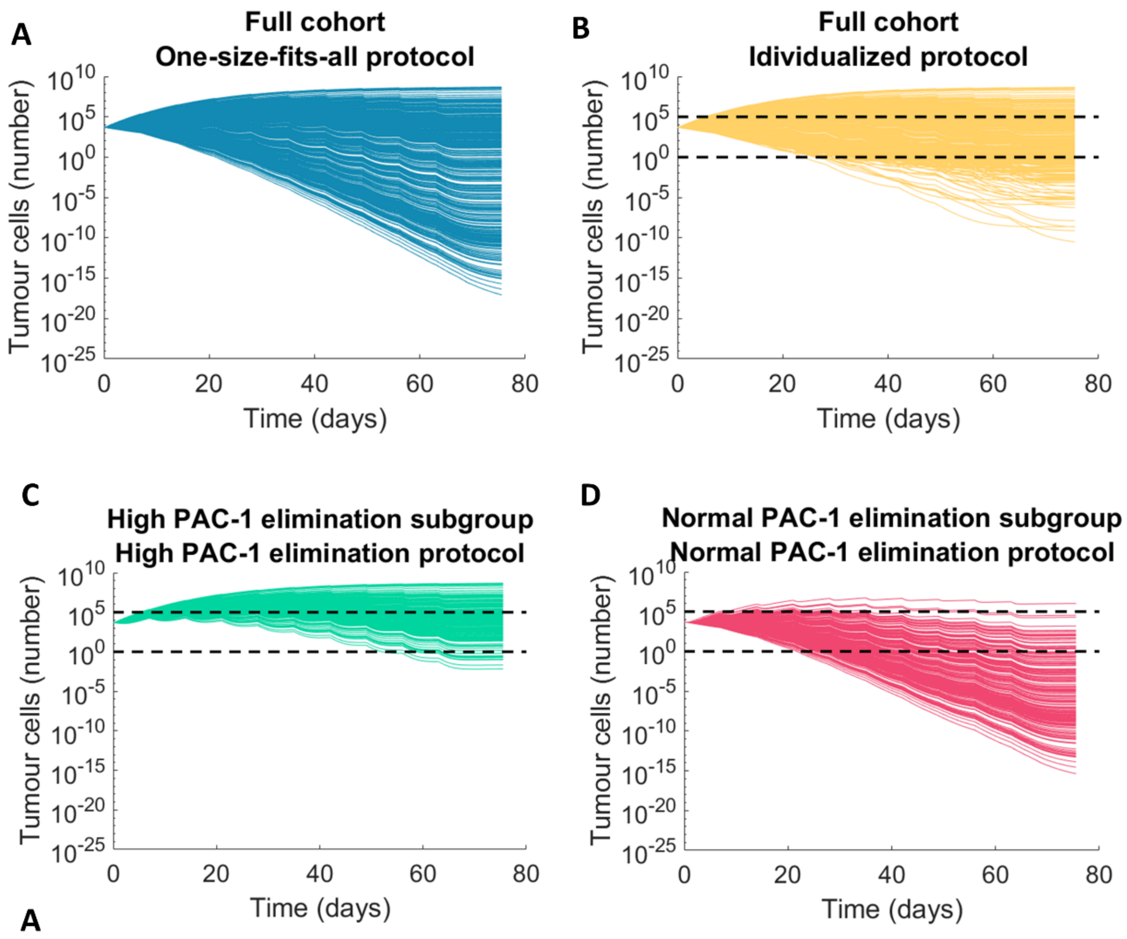


**Figure S7. Virtual cohort tumour growth under individualized, stratified and one-size-fits all protocols.** Virtual patient tumour cell numbers were simulated under the (A) full cohort level optimal protocol, (B) individualized protocols (C)-(D) subgroup protocols (High PAC-1 elimination subgroup treated with the High PAC-1 elimination protocol and Normal PAC-1 elimination subgroup treated with the Normal PAC-1 elimination protocol). Individual virtual patient trajectories are given on a log scale plot. The dotted lines indicate threshold markers for a number of tumour cells of 1 and $10^5$.

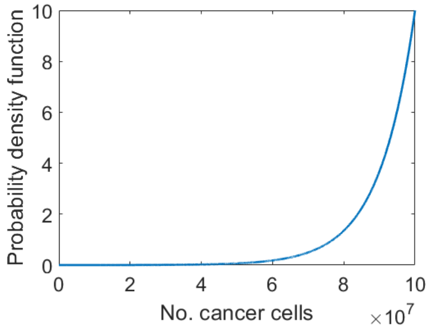


**Figure S8. Probability density function for virtual patient survival based on the number of cancer cells.** At each day, the patient was checked to see if they had survived by drawing a uniform random variable $U(0,1)$ and checking whether it was less than the probability of survival based on the number of cancer cells they had that day. Virtual patients were considered to not survive their treatment when their number of cancer cells exceeded $10^8$.
